# Vagus nerve stimulation limits the germinal center B cell response via CD4+ T cell-derived acetylcholine

**DOI:** 10.1101/2023.10.25.563818

**Authors:** Izumi Kurata-Sato, Ibrahim T Mughrabi, Minakshi Rana, Michael Gerber, Yousef Al-Abed, Barbara Sherry, Stavros Zanos, Betty Diamond

**Affiliations:** Center for Autoimmune Musculoskeletal and Hematopoietic Diseases, The Feinstein Institutes for Medical Research, Northwell Health, Manhasset, NY, USA; Institute of Bioelectronic Medicine, The Feinstein Institutes for Medical Research, Northwell Health, Manhasset, NY, USA; Donald & Barbara Zucker School of Medicine at Hofstra/Northwell, Manhasset, NY, USA; Department of Molecular Medicine, Donald and Barbara Zucker School of Medicine at Hofstra/Northwell, Hempstead, NY, USA; Center for Immunology and Inflammation, The Feinstein Institutes for Medical Research, Northwell Health, Manhasset, NY, USA

**Author notes:** These authors contributed equally (Izumi Kurata-Sato, Ibrahim T Mughrabi). Correspondence to Betty Diamond.

## Abstract

Neural signals are known to contribute to immune regulation and modulation of the innate immune response downstream of the vagus nerve has been well studied. The effects of vagus nerve activity on antibody production, however, have been largely unexplored. Here we use a chronic vagus nerve stimulation (VNS) mouse model to study the effect of vagal activation on T-dependent B cell responses. We observed lower titers of high-affinity serum IgG and fewer antigen-specific germinal center (GC) B cells in the spleen. GC B cells from chronic VNS mice expressed more active caspase-3 and exhibited an altered gene expression profile suggesting increased susceptibility to apoptosis and impaired maturation. Follicular dendritic cell (FDC) cluster dispersal and altered FDC gene expression suggested poor FDC function. These alterations were diminished in the absence of a subset of acetylcholine-producing CD4+ T cells. In vitro studies revealed that α7 and α9 nicotinic acetylcholine receptors (nAChRs) directly regulated B cell production of TNF, a cytokine crucial to FDC clustering. Engagement of the α4 nAChR subunit on B cells impaired Akt phosphorylation, presumably decreasing B cell survival. Thus, VNS-induced GC impairment can be attributed, at least in part, to the effect of acetylcholine on B cell intrinsic pathways, resulting in hindered B cell survival and maturation and leading to an alteration in FDC function. Our findings identify a potential therapeutic target to prevent immunosuppression in conditions associated with increased vagal activity.

## Introduction

Animal and clinical studies have demonstrated neural modulation of immunity ^1–4^. The mechanisms and pathways by which the nervous system controls immunity are intensively investigated ^5,6^, with the vagus nerve being the focus of many studies ^7–9^. Activation of the vagus nerve leads to activation of the splenic nerve which releases norepinephrine in the spleen. A specific subset of splenic CD4+ T cells that expresses choline acetyltransferase (ChAT) responds to norepinephrine with the release of acetylcholine (ACh) ^10^. Notably, ACh produced by CD4+ T cells exerts an immunosuppressive effect on splenic macrophages through engagement of α7 nicotinic acetylcholine receptors (nAChRs) ^6^. It is known that several nicotinic and muscarinic ACh receptors are expressed on the surface of lymphocytes also ^11^, but their immunomodulatory effects are less well studied.

The vagus nerve is activated in different inflammatory conditions, including sepsis ^12,13^. In the acute phase of sepsis, engagement of the vagus-immune pathway is beneficial as it suppresses potentially harmful inflammation to protect organ function ^14^. However, we have previously found that increased vagus nerve activation is sustained in post-sepsis mice, leading to the diminished responsiveness of splenic macrophages to LPS ^15^. We also reported an impaired germinal center (GC) response in post-sepsis mice ^16^. Decreased TNF production from antigen-activated B cells was associated with follicular dendritic cell (FDC) dispersion and lower titers of high-affinity antigen-specific antibody following immunization with a T-dependent (TD) antigen ^16^. Clinical studies also suggest insufficient humoral responses in sepsis survivors with poor vaccine responses and worse five-year survival rates due to infections compared to those who do not experience sepsis ^17,18^. These observations led us to hypothesize that vagus nerve stimulation alters the GC response.

The formation of the GC is a crucial process during the maturation of the antibody response. GCs are specialized microstructures formed in secondary lymphoid organs in response to antigen exposure. Following antigen activation, follicular (FO) B cells form GCs in cooperation with FDC clusters and T follicular helper (TFH) cells ^19^. FDCs present antigens as immune complexes ^20^, while TFH cells provide activation signals such as IL-21 and costimulatory molecules to GC B cells ^21^. In the GC, B cells undergo affinity-based selection ^22^ and mature into antibody-secreting plasma cells (PCs) and memory B cells. The modulation of the GC response by the vagus nerve has not been explored although B cells express α4β2, and α7 and α9 homomeric nAChRs ^11,23–26^.

Here we report that vagus nerve activation produces a reduced antibody response following immunization with a TD antigen. Mice subjected to chronic VNS produced a lower titer of high-affinity antibody. They also exhibited a lower number of antigen-specific GC B cells and decreased FDC clustering. The suppressive effect of chronic VNS was diminished upon removal of ChAT+ CD4+ T cells. Moreover, we demonstrate with *in vivo* and *in vitro* studies that ACh acts directly on B cells to reduce TNF production and Akt signaling, which are both critical for development of an effective GC response. These results demonstrate one underlying mechanism for B cell hyporesponsiveness in conditions of increased vagus nerve activation and suggest a potential therapeutic approach to modulating the adaptive immune response.

## Results

### Chronic vagus nerve stimulation reduces antigen-specific germinal center B cells

To assess the effects of VNS on the humoral immune response, we implanted mice with cuff electrodes on the left cervical vagus nerve, through which they received 5 min of VNS twice daily while freely behaving in their cages (Figure 1A). To assess longitudinal implant functionality and ensure successful engagement of the vagus nerve by stimulation, VNS-elicited reduction in heart rate was measured using implanted ECG electrodes (Figure S1A). Mice received chronic VNS for 14 days prior to immunization with the TD antigen, 4-hydroxy-3-nitrophenylacetyl (NP) chicken γ-globulin (CGG) (NP-CGG) in alum. VNS was continued for an additional 14 days to assess the effect on the antibody responses (Figure 1B). Sham mice underwent surgery but were not implanted with VNS electrodes.

**Figure 1.**
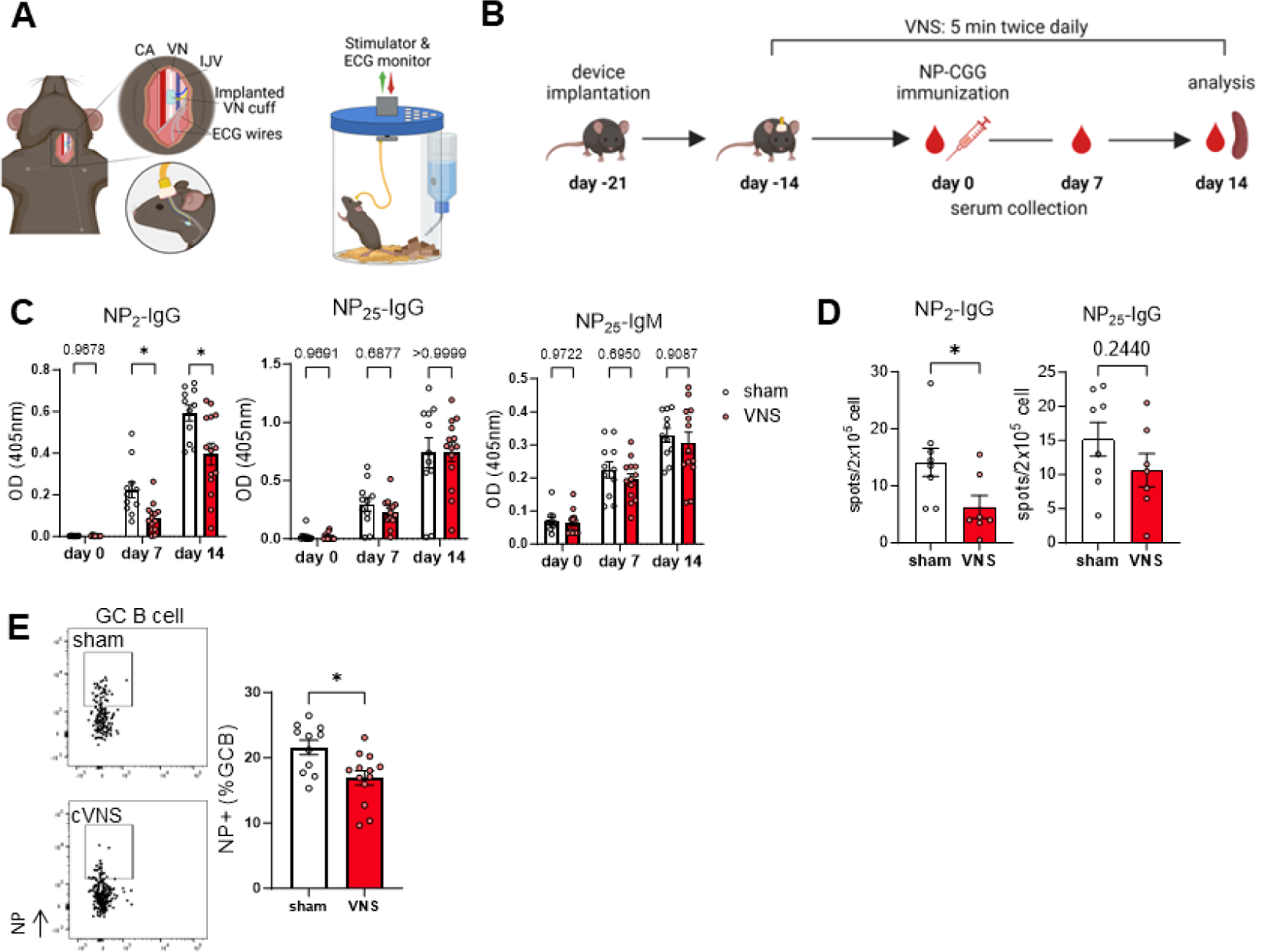
Chronic vagus nerve stimulation reduces the germinal center response. Mice implanted with a stimulation electrode on the left cervical vagus nerve received 5 min of VNS twice a day for 4 weeks while they were freely moving; a T-dependent B cell response was assessed in these animals. **(A)** Schema of chronic vagus implant and home cage with integrated VNS system. **(B)** Experimental schema of chronic VNS. **(C)** Quantification of high-affinity (NP_2_) and low-affinity (NP_25_) anti-NP antibody titers in serum. **(D)** ELISpot assays of anti-NP antibody secreting cells on day 14 post immunization. **(E)** NP-specific GC B cells in spleen on day 14. Representative flow cytometry plots for each condition. Data are shown as mean +/-SEM with each symbol representing an individual mouse. Data are combined from two independent experiments. Asterisks indicate significant differences (*p<0.05, **p<0.01, ***p<0.001, ns not significant) obtained using two-way ANOVA (C) and Mann-Whitney U test (D-E). CA: carotid artery, VN: vagus nerve, IJV: internal jugular vein. (A)(B); created with BioRender.com

Serum from chronic VNS mice showed decreased high-affinity (anti-NP_2_) IgG titers, whereas titers of low-affinity (anti-NP_25_) IgG were comparable to those of sham mice (Figure 1C). Total IgG was also comparable (Figure S1B). We enumerated NP-specific antibody-secreting cells in spleens by ELISpot assay and found a reduction of anti-NP_2_ IgG secreting cells in chronic VNS mice, but no difference in anti-NP_25_ IgG secreting cells (Figure 1D). Since high-affinity IgG plasma cells (PCs) mainly are derived from GCs, we next analyzed the frequency of NP-specific GC B cells. Chronic VNS decreased the frequency of NP-specific GC B cells (Figure 1E, gating strategy shown in Figure S1C). These data indicate that chronic VNS causes a diminished antigen-specific GC response with less high-affinity antibody production.

### Chronic VNS impedes GC B cell maturation and survival

GCs are composed of a light zone (LZ) and a dark zone (DZ), each characterized by distinct cellular dynamics and interactions ^19^. The LZ contains FDCs, TFH cells, and centrocytes where B cells undergo positive selection based on affinity for antigen. Centrocytes with weak antigen recognition undergo apoptosis whereas positively-selected centrocytes express CXCR4, become centroblasts, and move to DZ where they clonally expand ^27^. We sought to identify mechanisms underlying chronic VNS-induced GC impairment. We found an altered composition of GC B cells in chronic VNS mice with an increased percentage of centrocytes and a decreased percentage of centroblasts (Figure 2A). Both GC subsets had higher expression of active caspase-3, an early apoptosis marker (Figure 2B), suggesting diminished survival of GC B cells in chronic VNS mice. Furthermore, qPCR of GC B cells revealed increased PAX5, BCL6 and BACH2 and decreased PRDM1 (also known as Blimp1) mRNA expression in chronic VNS mice (Figure 2C). PAX5 maintains a B cell state by repressing expression of PC genes like XBP1. BCL6 and BACH2 collaborate to repress the transcriptional program associated with PC differentiation, including the expression of PRDM1 ^28,29^. To confirm these observations, we performed a single-cell transcriptome analysis of GC B cells. We identified five distinct GC B cell clusters among B220+ GL7+ NP+ splenic cells (gating strategy is shown in Figure S2A-D). Centrocytes were identified based on their relatively elevated expression levels of Cd83 and Cd86, and centroblasts were identified by high expression of Cxcr4 (Figure S2E). We observed a reduction in the proportion of proliferating centroblasts with heightened expression of histone genes, MKI67, and CDK1 (Figure 2E. Gene expression profiles are shown in Figure S2F, G). We observed a cluster of centrocytes that trended higher in chronic VNS mice (p= 0.1641), characterized by elevated expression of PAX5 and BCL6 mRNA and less XBP1 mRNA expression in comparison to the other centrocyte populations (Figure S2G), suggesting that these cells may be less exposed to differentiation signals provided by FDCs and TFH cells (Figure 2E). These findings indicate that GC B cells in chronic VNS mice exhibit a transcriptional program consistent with impaired survival and differentiation to PCs, which would contribute to the reduced NP-specific GC B cell response we observed.

**Figure 2.**
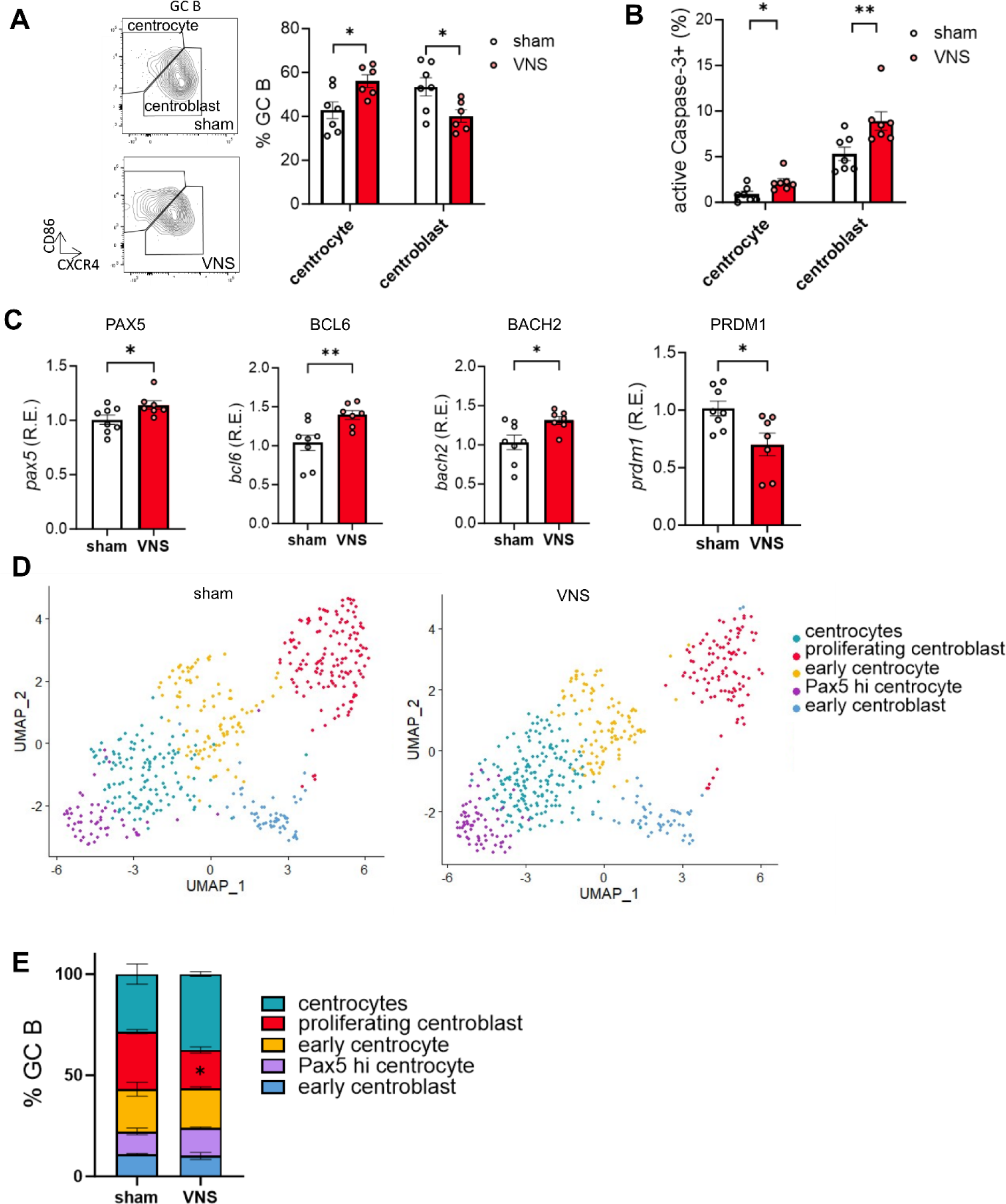
Alteration of germinal center B cell composition and survival. Splenic B cells were collected from VNS and sham mice on day 14 and analyzed. **(A)** Centrocytes (LZ GC B cells) and centroblasts (DZ GC B cells) in VNS mice. **(B)** Frequency of active caspase-3+ apoptotic cells in distinct GC B cell subsets assessed with flow cytometry. **(C)** Gene expression of sorted GC B cells was assessed with qRT-PCR. Relative expression to Polr2a is shown. **(D)** UMAP of GC B cell of sham and VNS mice. Each cluster was annotated using differentially expressed genes with the other clusters found by single-cell RNA seq (details shown in Supplementary figures). **(E)** Proportion of each GC B cell cluster. Data are shown as mean +/-SEM with each symbol representing an individual mouse from two (A-C) or single (D-E) independent experiments. Asterisks indicate significant differences (*p<0.05, **p<0.01, ns not significant) obtained using Mann-Whitney U test.

### Chronic VNS impairs follicular dendritic cell organization

FDCs cluster in B cell follicles and present antigens as immune complexes ^20,30^. Together with TFH cells, they provide signals to B cells to promote somatic hypermutation and differentiation into PCs ^31,32^. Maintenance of FDC clusters is known to be dependent on TNF, which is believed to originate from B cells ^30,33^. In chronic VNS mice, gene expression analysis revealed a reduction of TNF mRNA expression in GC B cells (Figure 3A), prompting us to examine FDC clusters. The relative abundance of FDCs identified as CD35+ CD45-was not different between chronic VNS and sham mice (Figure S3A, B).

**Figure 3.**
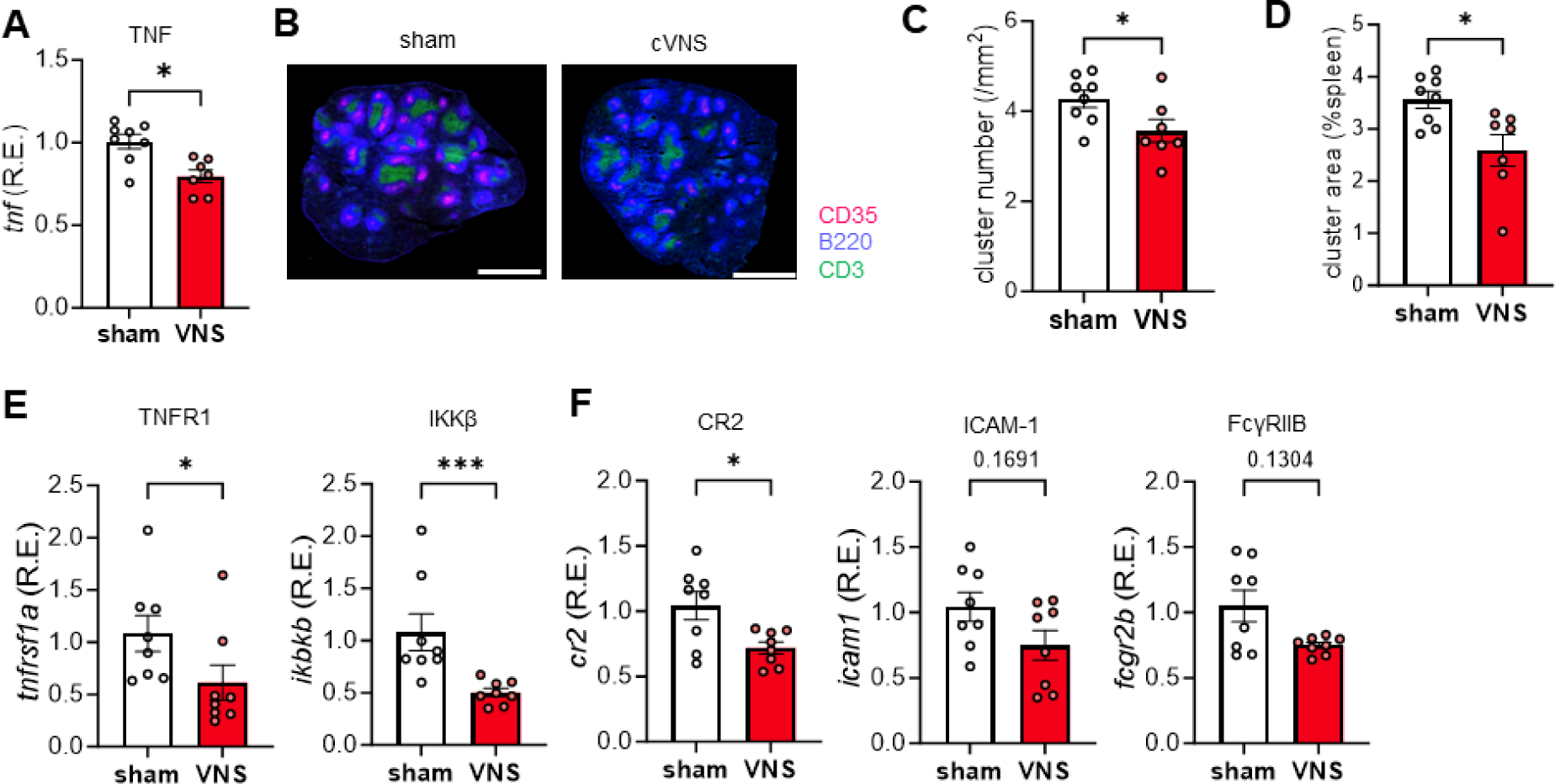
Chronic vagus nerve stimulation causes follicular dendritic cell dispersion, decreased TNF production by GC B cells and altered FDC gene expression. Spleens were harvested from chronic VNS and sham mice at day 14 and analyzed. **(A)** TNF gene expression in total GC B cells sorted from the spleens at day 14. The expression was assessed by qRT-PCR. **(B)** Representative image of chronic VNS and sham spleens stained with anti-CD35 (magenta), anti-B220 (blue) and anti-CD3 (green) antibodies. Magenta spots were analyzed as FDC clusters. White bar at bottom indicates 1,000 µm. FDC cluster numbers **(C)** and areas **(D)** were assessed. Two different slides per mouse were analyzed and averaged. Cluster numbers were counted manually. Areas were assessed with ImageJ software. **(E)(F)** Gene expression in FDCs sorted from chronic VNS and sham mice at day 14. Expression of the indicated genes was assessed by qRT-PCR. Data are shown as mean +/-SEM with each symbol representing individual mouse combined from two independent cohorts. Asterisks indicate significant differences (*p<0.05, **p<0.01, ****p<0.0001, ns not significant) obtained using Mann-Whitney U test.

Immunohistochemical analysis targeting CD35 demonstrated a decrease in the number and size of FDC clusters in the spleens of chronic VNS mice (Figure 3B-D), indicating a dispersal of FDCs. TNFR1 and downstream IKKβ mRNA expression were reduced in FDCs of chronic VNS mice (Figure 3E). TNFR1 is required for mature FDC networking. It activates a canonical NFκB pathway involving IKKβ/ NEMO complexes ^33,34^ that is indispensable for GC B cell maturation ^34^. Previous papers have shown that expression of the complement receptor, CR2, is crucial for immune complex trapping ^34,35^. The mRNA expression level of TNFR1, CR2 and IKKβ was reduced in FDCs of chronic VNS mice compared to sham mice; ICAM-1 and FcγRIIB mRNA levels showed a trend toward decreased expression (Figure 3F). These observations suggest diminished functionality in FDCs of chronic VNS mice.

### ChAT+ CD4+T cell mediates GC impairment

To address a potential cellular mediator of the altered B cell response in chronic VNS mice, we examined the impact on ChAT+ T cells. ChAT reporter mice revealed an increase in ChAT+ CD4+ T cells in the spleen of chronic VNS mice (Figure 4A). Approximately 30% of ChAT+ CD4+ cells had a TFH phenotype (CXCR5+, PD-1+) while only 5% of ChAT-CD4+ cells were TFH-like (Figure S4). We also studied mice lacking ChAT+ T cells to address the impact of ChAT+ T cells on the GC response. Chronic VNS mice lacking ChAT+ CD4+ T cells had more serum anti-NP_2_ IgG and an increased frequency of NP-specific GC B cell compared to control (CD4-cre ChAT^wt^) chronic VNS mice (Figure 4B-D). Importantly, FDC clusters were enlarged in these mice; the number of clusters did not differ between mice with or without ChAT+ T cells (Figure 4E-G). These data indicate that ChAT+ CD4+ T cells attenuate the GC response in chronic VNS mice.

**Figure 4.**
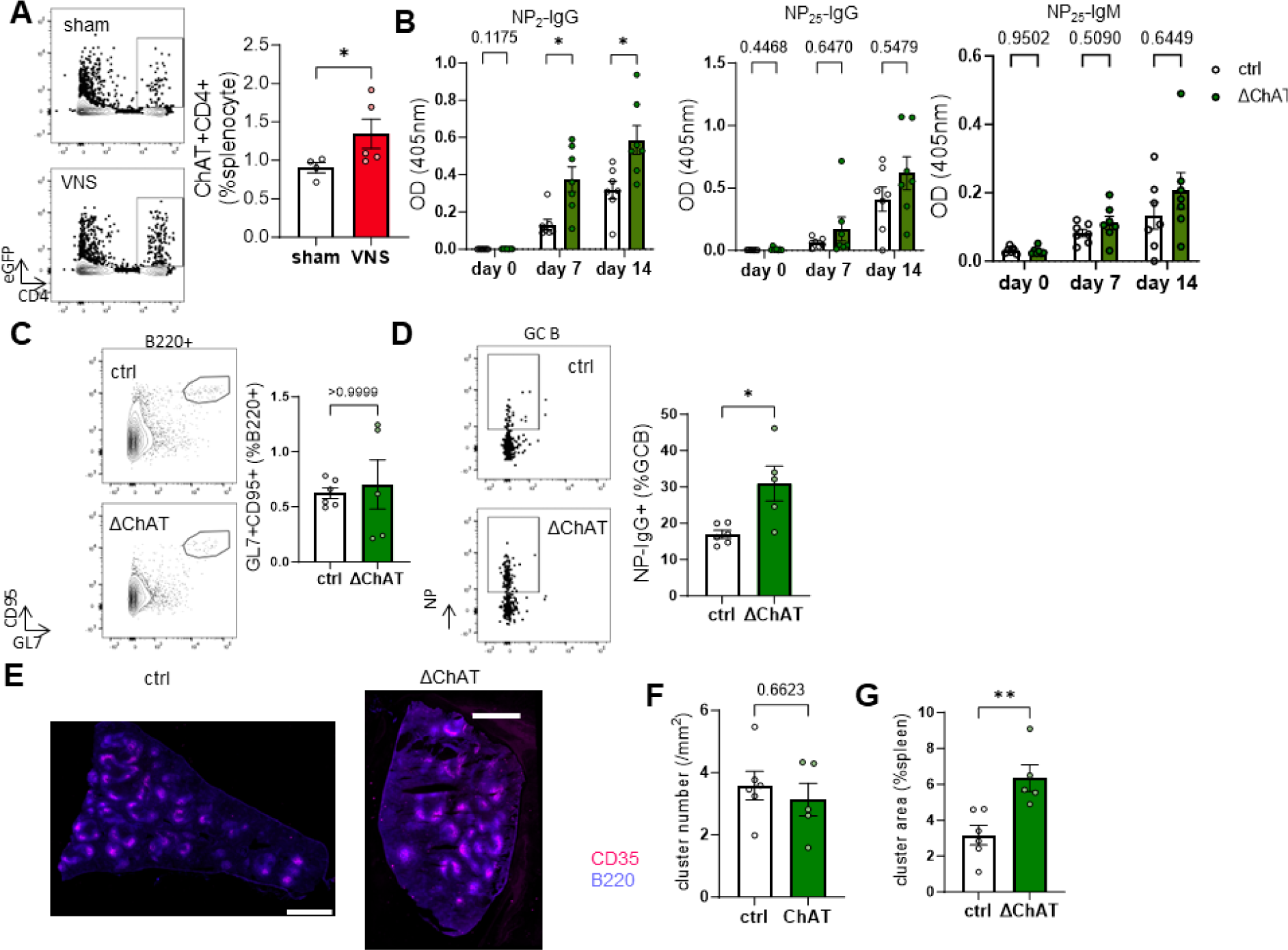
CD4+ ChAT+ T cells impair the germinal center B cell response during chronic vagus nerve stimulation. **(A)** Frequency of ChAT+ T cells was analyzed using ChAT^eGFP^ reporter mice. Chronic VNS and sham mice were analyzed on day 14. **(B)** NP-specific antibody titers in serum from CD4-cre ChAT^WT^ (control) and CD4-cre ChAT ^flox/flox^ (ΔChAT) mice. Total **(C)** and NP-specific **(D)** GC B cells in spleens on day 14. **(E)** Representative images of splenic FDC clusters in control and ΔChAT mice. The numbers **(F)** and areas **(G)** were assessed using the same method as Figure 3B. Data are shown as mean +/-SEM with each symbol representing an individual mouse from one (A) or two (B-D) independent cohorts. Asterisks indicate significant differences (*p<0.05, ns not significant) obtained using two-way ANOVA (B) or Mann-Whitney U test (A, C, D, F, G).

### AChR engagement directly impairs signaling pathways in B cells

Given our findings that implicate ChAT+ CD4+ T cells in the effects of chronic VNS on the GC response, we explored the direct impact of ACh on B cells. FO B cells stimulated *in vitro* with CpG, a toll-like receptor (TLR) 9 agonist, increased transcription of TNF. Increasing doses of nicotine, a pan-nicotinic ACh receptor (nAChR) agonist, decreased CpG-induced TNF mRNA in a dose-dependent fashion (Figure 5A, the gating strategy of FO B cell is described in Figure S5A). Immunofluorescence showed decreased NFκB p65 subunit translocation to the nucleus (Figure 5B), a mechanism known to result in decreased TNF production ^36^.

**Figure 5.**
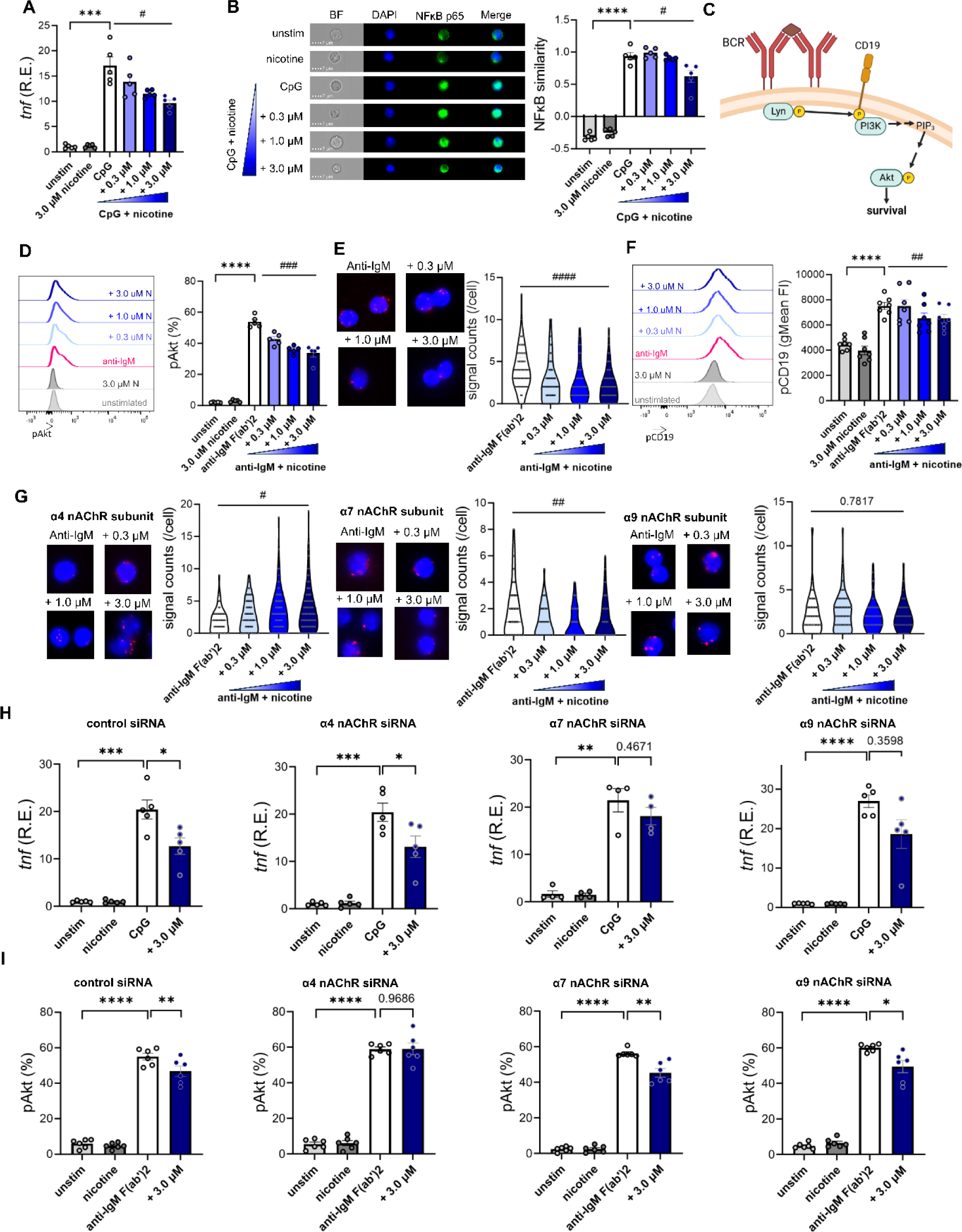
Acetylcholine directly regulates TLR9 signaling and BCR-mediated Akt phosphorylation via nicotinic receptors. **(A)** TNF mRNA in nicotine exposed, CpG-treated FO B cells. Splenic FO B cells from untreated mice were sorted using flow cytometry and incubated with 2.5µM CpG and/or the indicated amount of nicotine. Cells were collected after 4 hours incubation and qRT-PCR was performed with the extracted mRNA. **(B)** NFκB p65 translocation to nucleus under CpG with or without nicotine stimulation. Total B cells were isolated from untreated mice using magnetic sorting. Cells were incubated with the indicated amount of nicotine and/or 1.0 µM CpG for an hour and fixed. NFκB p65 translocation was assessed by comparing similarity of DAPI (blue) and NFκB (green) staining morphology using imaging cytometry. **(C)** Scheme of BCR signaling pathways. Created with BioRender.com. **(D)** Phosphorylation of Akt (S473) following BCR activation with or without nicotine exposure. Splenocytes were collected from untreated mice and stimulated with or without nicotine and anti-IgM F(ab)’2. Phosphorylated Akt in FO B cells was quantified using flow cytometry. **(E)** Proximity ligation assay with membrane IgM and CD19. Total B cells were isolated from untreated mice and incubated with anti-IgM F(ab)’2 with or without nicotine. Proximity of membrane IgM and CD19 was assessed by counting signal spots. Each dot indicates cell. Three independent fields were analyzed per sample. **(F)** Phosphorylation of CD19 with BCR activation and nicotine exposure assessed using the same method as (D). **(G)** Proximity ligation assay with membrane IgM and nicotinic ACh receptor subunits, assessed with the same method as (F). **(H)** TNF mRNA in nAChR-knocked down FO B cells. B cells were transfected with siRNA for each nAChR subunit or control and subsequently stimulated with CpG and nicotine. **(I)** Akt (S473) phosphorylation of nAChR-knocked down FO B cells. B cells were transfected with siRNA for each nAChR siRNA and subsequently incubateded with anti-IgM F(ab’)2 and nicotine. Data are shown as mean +/-SEM (A, B, D, F, H, I). Data from two to three independent experiments were combined and assessed. Hash marks indicate significant differences (^#^p< 0.05, ^##^p< 0.01, ^###^p< 0.001, ^####^p< 0.0001) obtained using one-way repeated measure ANOVA test with Geisser-Greenhouse correction. Asterisks indicate significant differences (*p<0.05, **p<0.01, ***p<0.001, ****p<0.0001) obtained using paired T test. N: nicotine.

We next examined phosphorylation of Akt downstream of BCR crosslinking (Figure 5C), which is essential for antigen-mediated B cell survival ^37,38^. Akt phosphorylation was specifically inhibited by nicotine (Figure 5D). As Akt activation requires proximity of the BCR with CD19 ^39^, we performed a proximity ligation assay and found that nicotine blocked the coligation of CD19 to the BCR (Figure 5E) and subsequent CD19 phosphorylation (Figure 5F) without altering CD19 and BCR expression (Figure S5B, C). The expression of FcγRIIB and downstream SHIP1 activation, which are known to regulate the PI3K-Akt pathway ^40^, were not altered (Figure S5D, E). We hypothesized that nAChRs might physically interact with BCRs upon ACh engagement and prevent CD19 from entering the lipid raft where BCRs are located. Proximity ligation analysis showed that nAChRs with α4 subunits translocated toward BCRs in response to nicotine binding (Figure 5G).

To understand further the role of each nAChR, we performed siRNA-mediated knock down of specific nAChR subunits (Figure S5F, G). The inhibition of TNF production was abolished in α7 and α9 nAChR-KD cells but not in α4 nAChR-KD cells (Figure 5H). In contrast, inhibition of Akt phosphorylation was diminished in α4 nAChR-KD cells (Figure 5I). We additionally used cells from B cell-specific α7 nAChR KO mice to confirm that α7 nAChR is important for the inhibition of TNF production; there was no effect on Akt phosphprylation in α7 nAchR cKO mice (Figure S5H, I). These data suggest ACh directly alters B cell activation and function through distinct nAChR-mediated pathways.

## Discussion

In the present study, we studied the effects of chronic VNS on the B cell response to a TD antigen. We demonstrated that chronic VNS leads to a reduction in the high-affinity antibody response with impaired GC formation and fewer antigen-specific GC B cells. GC B cells in chronic VNS mice exhibited an increased propensity to undergo apoptosis and impaired maturation into PCs. FDCs were dispersed and had an altered gene expression profile. In the presence of nicotine, B cells stimulated with CpG secreted less TNF and B cells stimulated through the BCR exhibited less phosphorylation of Akt. Distinct nAChRs were responsible for these effects. Taken together, our data indicate that the poor GC response in chronic VNS mice results from at least two distinct mechanisms: decreased TNF release from B cells triggering functional alterations in FDCs and impaired formation of the BCR-CD19 coreceptor complex leading to reduced Akt signaling and B cell survival.

Few studies have investigated neural modulation of humoral immune responses. B cells are known to express nAChRs ^11^. The α7 nAChR has been documented to exert inhibitory effects on NFκB translocation to the nucleus and TNF release from macrophages, especially in the context of endotoxemia ^7,14^. Whether α7 nAChRs on macrophages regulate an endosomal TLR pathway, as we observed in B cells, has not been investigated.

Consistent with our findings, the α4β2 nAChR is reported to be located near the BCR ^23^. It is also important to note that TM4, a transmembrane domain of the nAChR, is thought to be a lipid sensor and influence the surrounding membrane lipid organization, particularly the lipid raft ^41^. It is plausible therefore to speculate that ACh may regulate the BCR complex through effects on lipid rafts after engagement of the BCR. Consistent with our findings, it is also previously reported that α7 and α9 nAChRs are not involved in BCR-mediated activation ^23^. In contrast to these findings, Zhang et al. reported that α9 nAChRs contribute to post-GC PC expansion ^24^. Their findings indicate that the splenic nerve promotes PC expansion through α9 nAChRs without affecting the GC B cell population. A distinction between the two studies lies in our approach of applying chronic VNS 14 days before immunization, leading to FDC dispersion at the time of immunization which was not the case in their study. Also, we adopted a regimen of twice-daily, 5-minute VNS according to our previous finding that this regimen reduced serum TNF levels in response to LPS ^42^. Our regimen suppressed the GC response, but it is possible that distinct VNS parameters act differently on humoral immunity. In the present study, we found that different nAChRs play distinct roles. Given the varying binding affinities of these receptors ^43^, it is plausible that the outcome of ACh exposure differs based on the amount of ACh released.

The findings on FDC dispersion align with our previous findings in post-sepsis mice in which the vagus nerve is constitutively activated ^15,16^, suggesting the vagus nerve plays a role in impairing FDCs and thus dampening humoral immunity. However, it is essential to acknowledge that chronic VNS, as delivered in this study, did not fully replicate the post-sepsis condition. Post-sepsis mice displayed a more pronounced suppression in antibody production, affecting both high and low affinity antibodies ^16^. This may indicate the involvement of additional mechanisms or heightened vagus nerve activity.

It is also important to acknowledge that there could be other neurotransmitters and cell types at play during chronic VNS. Norepinephrine and dopamine are also reported to be increased by VNS ^44^ and immune cells including B cells possess receptors for these molecules ^45–49^. As mice with ChAT deficient CD4+ T cells showed only a partial abrogation of the VNS effect, other pathways likely play a role in our model. Of interest, Suzuki et al. showed β2 adrenergic receptors prevented T and B cell egress from lymph nodes through physical interactions with chemokine receptors ^50^. Furthermore, TFH cells may be involved in VNS-mediated changes in GC response because CD4+ T cells also express both nAChRs and adrenergic receptors ^11^. We found an increase of CXCR5 expression of ChAT+ CD4+ T cells in chronic VNS mice (Figure S4A), suggesting chronic VNS may affect migration of these cells within the spleen, ultimately contributing to modulation of the GC response.

Stimulation of the cervical vagus nerve, where all afferent and efferent vagal fibers converge, engages several autonomic circuits and reflexes ^6^, whose activation was not evaluated in this study. Some of these pathways, including pre-ganglionic, sympathetic fibers in the splanchnic nerve that project to the celiac ganglion ^6^, may be involved in the effects of chronic VNS that we report. Their contribution will have to be determined in future studies.

In summary, we have revealed novel pathways explaining how the vagus nerve helps control humoral immunity. These findings have potential clinical significance in reversing undesired immunosuppression, which is seen in sepsis survivors ^17,18,51^. Altering vagus nerve activity could be a new therapeutic strategy to reverse this immunosuppressed state which may be present also in survivors of severe COVID-19 infection.

## Acknowledgements

The authors thank Flow Cytometry Core Facility and Microscopy Core Facility of the Feinstein Institutes for Medical Research for technical assistance. The authors also thank Houman Khalili for assistance on performing single-cell RNA seq. This work is supported by Uehara Memorial Foundation Overseas Research Fellowship to IKS.

## Author contributions

IKS and ITM designed and conducted experiments, analyzed data, and wrote the manuscript. MR designed experiments. MG conducted experiments and analyzed data. YA, SZ, and BS conceptualized the study and designed experiments. BD conceptualized the study, designed experiments, interpreted data, and wrote the manuscript.

## Materials and methods

### Mice and immunization

Male C57BL/6 were purchased from The Jackson Laboratory and Charles River Laboratories and used at 8-15 weeks of age. ChAT^eGFP^ mice were kind gift from Dr. Kevin Tracey. CD4-cre, ChAT^flox^, CD19-cre and CHRNA7^flox^ mice were purchased from Jackson Laboratory and bred at the Feinstein Institutes for Medical Research for more than 10 generations (detailed information of strains is found in Table S1). All mice were housed under specific pathogen-free conditions. In some experiments, mice were administered 100 μg of NP-CGG (Biosearch Technologies) emulsified in 100 μL of Imject^TM^ alum (Thermo Fisher Scientific) by intraperitoneal injection. Mice were euthanized by CO_2_ overdose or by cervical dislocation at the time points indicated in the Results. Serum was obtained by tail bleeding or by cardiac puncture. All protocols including vagus nerve stimulation were approved by the Institutional Animal Care and Use Committee.

### Vagus nerve and ECG electrodes implantation

Commercial micro-cuff electrodes (100 µm microsling, CorTec, Germany) and platinum iridium wires were integrated onto a nano-strip connector (Omnetics). The cuff electrodes were implanted on the left cervical vagus nerve and the platinum iridium wires were tethered subcutaneously to the chest wall to record ECG as described previously ^42^. In brief, anesthesia was induced with 3-4% isoflurane and maintained with 1.5%. On a warmed surgical pad, an area over the dorsal skull was exposed and scrubbed with saline and hydrogen peroxide and a subcutaneous tunnel to the ventral neck was created. The carotid sheath was exposed through a ventral neck incision and the left vagus nerve was gently dissected away from the sheath. The cuff and ECG electrodes were tunneled to the ventral neck through the skull incision and the cuff electrode was carefully placed on the vagus nerve and embedded under neck musculature. The ECG electrodes were sutured subcutaneously to the upper and lower chest wall through the ventral neck and a left subcostal incision. The Strip connector was then cemented to the skull and sealed with surgical glue. The surgery was performed under strict aseptic technique. Mice were supplemented with saline throughout the procedure and treated with meloxicam (5 mg/kg) post-operatively. Mice were allowed to recover for about a week before starting chronic VNS.

### Chronic vagus nerve stimulation

Mice were singly housed in custom-made cages (Fig 1A) each comprising a clear acrylic cylinder (6 inches diameter, 8 inches height), a 3D-printed ventilated plastic cover with an integrated commutator system (P1 Technologies), and a water bottle. Food and nestlet material were provided on the floor of the cage and were changed weekly. Mice were acclimated to these cages for one week prior to receiving the implant surgery and continued to be housed in their home cages till the end of the experiment. After one week of recovery, mice were connected to the commutator via a flexible cable which allows free movement. The commutator system was connected to a battery-operated stimulating/recording system (Neurochip 3 ^52^) located outside the cage and programmed with stimulation parameters and schedule. The custom-made cages were placed on a mobile station and housed in the mouse vivarium on a 12-h light/dark cycle with ad-libitum access to food and water. Mice received twice-daily stimulation (6 hours apart) using the following parameters: 500 µs pulse width, 30 Hz frequency, and 5 min duration at a current intensity that caused ∼20-30% reduction in baseline heart rate. The recorded ECG was reviewed daily to monitor the functionality of the stimulating electrode and adjust stimulation intensity to maintain a consistent heart rate response throughout the period of VNS. Mice were monitored daily to observe indicators of wellbeing, including food and water consumption and nest building. On weeks 0 and 1 of VNS (Figure 1B), blood was collected by tail-tip sampling under isoflurane anesthesia for serological assays. At the end of week 4 of VNS mice were sacrificed and spleen and blood collected for analysis.

### ELISA

Serum titers of NP-specific antibodies, total IgG and IgM were measured by ELISA. Half-area flat-bottom plates were coated with 10 µg/mL of NP_2_ or NP_25_ -BSA (Biosearch Technologies) or 5 µg/mL of goat anti-mouse IgG/IgM (Southern Biotech) overnight at 4℃. Plates were then washed 6 times with 0.05% Tween/PBS solution and blocked with 1% BSA/PBS at room temperature for an hour. Serum diluted with 0.2% BSA/PBS (1:2000 for anti-NP IgG, 1:20000 for total IgG/IgM) was added to the wells. After 2 hours of incubation, plates were washed and alkaline phosphatase (AP)-conjugated goat polyclonal anti-mouse IgG/IgM (Southern Biotech) diluted with 0.2% BSA/PBS (1:1000) was added and incubated for an hour, followed by color development with 1 mg/mL phosphatase substrate (Millipore Sigma). The color absorbance of each well was read at 405 nm on 1430 Multilabel Counter Spectrometer (Perkin Elmer).

### ELISpot

For counting NP-specific antibody producing cells, 96-well Immulon 4HBX plates (Thermo Fisher Scientific) were coated with 10 µg/mL of NP_2_- or NP_25_-BSA in PBS overnight at 4℃. Plates were washed and blocked with RPMI1640 (Gibco) supplemented with 10% heat-inactivated FBS, 1% Penicillin/Streptomycin and 10 mM HEPES for an hour at 37℃ to avoid nonspecific binding. Serially diluted splenocytes suspended in the supplemented RPMI1640 were added to the wells. Plates were incubated overnight at 37℃ and washed in the following day. Wells were incubated with biotinylated goat anti-mouse IgG/IgM (Southern Biotec) for 2 hours at 37℃, followed by AP-conjugated streptavidin (Southern Biotec) for an hour at 37℃. Spot development was performed using Vector Blue Alkaline Phosphatase Substrate Kit (Vector Laboratories) according to manufacturer’s instruction. Spots were counted manually.

### Flow cytometry and cell sorting

A single-cell suspension of spleens was obtained by mechanical dissociation. Cells were suspended in flow cytometry buffer (2% FBS, 0.1% NaN3, 1 mM EDTA in HBSS) at 1 x 10^7^ cells/mL and incubated with Fc block (anti-mouse CD16/32; clone 93. BioLegend) for 10 minutes before incubation with specific antibodies listed in Table S1, except for FcγRIIB staining. Dead cells were stained out with Fixable Viability Dye (Thermo Fisher Scientific). For detection of NP-specific B cells, biotinylated NP-BSA (Biosearch Technologies) was incubated with APC-conjugated streptavidin (BioLegend) before being added to the cell suspension. For active caspase-3 staining, Cytofix/Cytoperm buffer (BD) was used according to the manufacturer’s instructions. Cells were analyzed with a BD LSR Fortessa X-20 flow cytometer (Becton Dickinson) and Flowjo software v10.7.2 (Tree Star) or sorted with a FACS Aria (BD Biosciences). Sorted cells then underwent stimulation or were stored in TRIzol reagent (Life Technologies) at -80^0^C for mRNA extraction.

### Quantitative real-time PCR

Messenger RNA was extracted from cells using Direct-zol RNA Microprep kit (Zymo Research) according to the manufacturer’s instructions. Freshly isolated mRNA was reverse transcribed into complementary DNA with random primers by iScript cDNA Synthesis Kit (Bio-Rad).

Quantitative RT-PCR was performed on LightCycler 480 II (Roche) using TaqMan gene expression assay (Thermo Fisher Scientific). Primers used for assays are listed in Table S1. Relative gene expression was normalized to expression of RNA polymerase II subunit A (polr2a) using the 2^-ΔΔCt^ method. For GC B cell and FDC gene expression, synthesized cDNA was subjected to pre-amplification of target genes as well as polr2a using TaqMan PreAmp Master Mix Kit (Thermo Fisher Scientific) prior to qRT-PCR.

### Single-cell RNA seq

B220+GL7+NP+ cells were sorted from chronic VNS (n= 4) and sham (n= 4) mice at day 14 by flow cytometry. The gating strategy is described in Figure S2A. Each sample was tagged with different TotalSeq-C hashing antibody (Biolegend). Library preparation was performed according to the manufacturer’s instructions (Next GEM Single Cell 5’GEM v3.1 protocol, 10x Genomics). Libraries were sequenced by MedGenome on an Illumina NovaSeq. FASTQ data from 10x Chromium were processed with CellRanger v6.1.238 aligning to the C57BL/6 reference genome. The count matrix was loaded into R and further processed using the Seurat R package v4.9.9. Doublets and empty droplets were determined and excluded by analyzing hashing oligos. Cells with fewer than 200 distinct genes or with more than 5% of mitochondrial genes were also excluded as low-quality cells. The remaining cells were clustered using UMAP (resolution= 0.8). Four clusters which highly expressed *Cd3e* or *Itgam* and did not express *Ms4a1* were annotated as contamination of T and myeloid cells and subsequently excluded. Four clusters (3, 4, 6, 7) were determined as the GC B cell clusters using known markers (*Aicda*, *S1pr2*, *Fas* and *Bcl6*: Figure S2C). One chronic VNS and one sham mice were omitted due to relatively small number of GC B cells (< 50 cells). These clusters were extracted and re-analyzed for differential gene expressions (Figure S2D) and frequencies.

### Immunofluorescence

Freshly isolated spleens were half-cut, embedded in OCT compound (Fisher Scientific) and frozen at -80 ℃. OCT-embedded 8-µm cryostat sections were fixed with cold acetone for 10 min. Sections were then blocked in PBS with 8% normal rat serum for 1 hour prior to staining. Slides were stained with biotinylated anti-mouse CD35 (diluted in 1:100) (8C12, BD Bioscience), APC conjugated anti-mouse CD3 (1:100) (145-2C11, BD Bioscience) and Brilliant Violet 421 conjugated anti-mouse B220 (1:200) (RA3-6B2, BioLegend) in 2% normal rat serum/PBS overnight at 4 ℃. Slides were subsequently washed and incubated with Alexa Flour 568 conjugated streptavidin for 45 min at room temperature, followed by mounting with Dako Fluorescence Mounting Medium (Dako). Slides were analyzed with Zeiss Apotome 3 microscope (Zeiss). Imaging data were analyzed with ImageJ. To evaluate the FDC clusters, two independent slides were assessed and averaged per spleen.

For imaging cytometry, stimulated cells were fixed and permeabilized overnight with Foxp3/Transcription Factor Staining Buffer Set. Anti-NF-κB p65 antibody (diluted in 1:500) (D14E12, Cell Signaling Technology) was added and incubated for an hour, followed by AF488-conjugated anti-rabbit IgG. After washing, cells were stained with DAPI. NF-κB translocation was assessed with Amnis ImageStreamX Mk II (Luminex). For each condition 400 cells per mouse were collected and analyzed. The similarity score was obtained from IDEAS software, which calculated fluorochrome distribution similarity between anti-NF-κB antibody and DAPI.

### *In vitro* cell stimulation

For measurements of TNF mRNA, FO B cells were sorted (the gating strategy is shown in Figure S4A) and stimulated with 2.5 µM CpG (InvivoGen) and nicotine (Sigma) in RPMI1640 (Gibco) supplemented with 10% heat-inactivated FBS, 1% Penicillin/Streptomycin and 10 mM HEPES. After 4 hours of incubation, cells were collected and pellets were solubilized in TRIZOL reagent. These solution were stored at -80℃.

For NF-κB translocation assays, B cells were isolated from naïve mouse spleens with EasySep^TM^ Mouse Pan-B cell Isolation Kit (Stem Cell Technologies) and suspended at 1 x 10^6^ cells/mL in the supplemented RPMI1640 (Gibco). Cells were stimulated with 1 µM CpG with or without nicotine for an hour. After washing cells were fixed and permeabilized using Foxp3/Transcription Factor Staining Buffer Set.

### B cell receptor signal analysis

Single cell suspension were obtained from naïve mouse spleens and rested in the supplemented RPMI1640 at 37 ℃ for an hour before stimulation. Cells were then stimulated with nicotine for 20 min and subsequently anti-mouse IgM F(ab’)2 (Jackson ImmunoResearch) for 2 min (pCD19) or 5 min (pAkt). Stimulation was ceased by adding pre-warmed Lyse/Fix buffer (BD). Fixed cells were then incubated with Perm/Wash buffer (BD) for 30-60 min and stained with antibodies listed in Table S1.

### Western blot

Western blot of pSHIP1 and SHIP1 was performed as we previously described ^53^. Briefly, FO B cells were sorted and stimulated for 10 min following the same method described above. Whole anti-mouse IgM (Jackson ImmunoResearch) was used instead of anti-IgM F(ab’)2 to activate FcγRIIB. Stimulation was immediately ceased by adding ice-cold PBS. Cells were lysed in Cell Lysis buffer (BD) supplemented with Halt Protease and Phosphatase Inhibitor Cocktail (Thermo Fisher Scientific) at 1 x 10^5^ cells/3 µL. Capillary western blot was performed with Jess (Protein Simple) according to the manufacturer’s instruction. Obtained data were analyzed with Compass for SW software (Protein Simple). AUC of band intensity histograms was used for protein amount quantification. Antibodies used are listed in Table S1.

### Proximity ligation assay

B cells were isolated from naïve mouse spleens with EasySep^TM^ Mouse Pan-B cell Isolation Kit (Stem Cell Technologies) and stimulated following the same method as BCR signal analyses described above. The proximity of the interested molecules was visualized using Duolink In Situ Detection reagents (Millipore Sigma) according to the manufacturer’s instructions. Briefly, fixed cells were blocked for 50 min at 37 ℃ and labelled with antibodies listed in Table S1 for 30 min at room temperature in 96 well V-bottom plates. After a washing step, PLA probes were added in wells and incubated for an hour at 37 ℃. Cells were subsequently washed and incubated with ligase for 30 min at 37 ℃, followed by oligonucleotide amplification with DNA polymerase for 100 min at 37 ℃. Cell suspensions were applied to slide glasses using SHANDON Cytospin 4 (Thermo Scientific) and observed with Zeiss Apoptome 3 microscope (Zeiss).

### siRNA transfection

B cells were isolated from naïve mouse spleens with EasySep^TM^ Mouse Pan-B cell Isolation Kit (Stem Cell Technologies) and suspended in Opti-MEM™ I Reduced Serum Medium (Gibco). Each nAChR siRNA or control siRNA was co-transfected with BLOCK-iT™ Alexa Fluor Red Fluorescent Control (Thermo Fisher Scientific) to the cells using Lipofectamine RNAiMAX Reagent (Thermo Fisher Scientific), according to the manufacturer’s instruction. Alexa Fluor 555 positive cells were sorted out by flow cytometry after 4 hours of incubation. Cells underwent further assays after additional 20 hours of culture in RPMI1640 medium supplemented with 10% heat-inactivated FBS and 10 mM HEPES. siRNAs used are listed in Table S1.

### Statistical analysis

Statistical analysis was performed using Prism version 10 (GraphPad). P values <0.05 were considered statistically significant. The statistical method used in each experiment is reported in the figure legends.

