## supplementary figures and table for "Vagus nerve stimulation limits the germinal center B cell response via CD4+ T cell-derived acetylcholine"

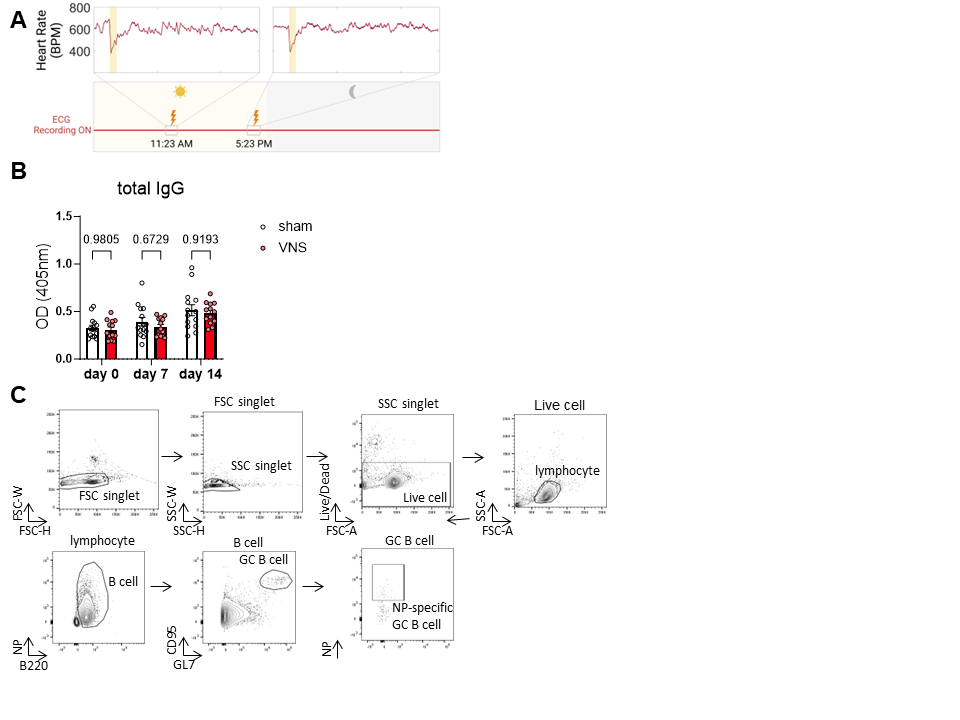

### Figure S1.

**(A)** Schema of ECG monitoring during chronic VNS. ECG was recorded and reviewed daily to monitor the functionality of the electrode throughout the period of VNS. This figure was created with BioRender.com. **(B)** Quantification of total IgG and IgM titers in sera. **(C)** The gating strategy of total and NP-specific GC B cells. Data are shown as mean +/- SEM with each symbol representing an individual mouse from two independent experiments. Two-way ANOVA was used to test significance.

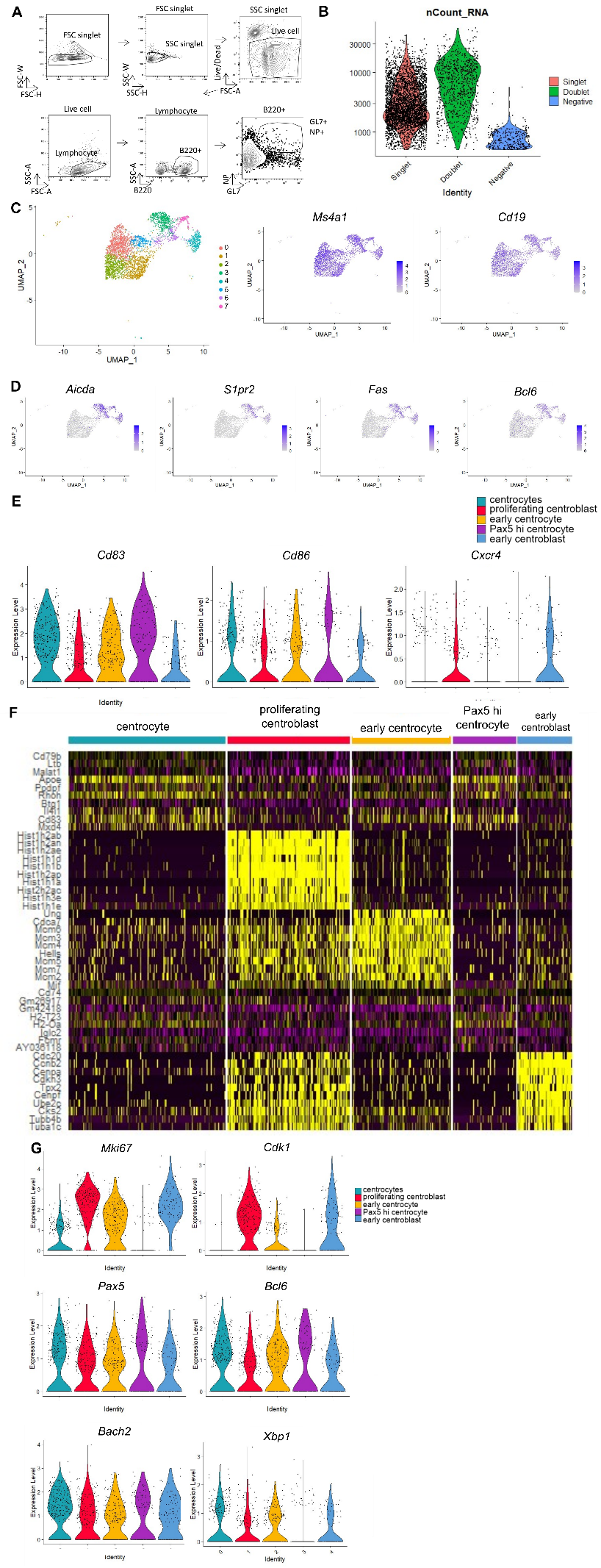

### Figure S2.

**(A)** The gating strategy of B220+ GL7+ NP+ cells. **(B)** Identification of singlet gel beads emulsions (GEMs) using hashing oligos. Doublets and empty (negative) GEMs were excluded from further analysis. **(C)** UMAP plot of total B cells. Eight clusters were identified. Feature plots of *Ms4a1* and *Cd19* verified the cell identity. **(D)** Feature plots of GC B cell marker genes (*Aicda*, *S1pr2*, *Fas* and *Bcl6*). Cluster 3, 4, 6, 7 were identified as GC B cell clusters. The four clusters were extracted and re-clustered. **(E)** Violin plots of *Cd83*, *Cd86* and *Cxcr4* among the re-analyzed GC B cell clusters. These parameters were used to annotate centrocytes and centroblasts. **(F)** Gene expression heatmap of the GC B cell clusters using top 10 differentially expressed genes of each cluster. **(G)** Violin plots of proliferation and B cell maturation genes. CC: centrocyte, CB: centroblast.

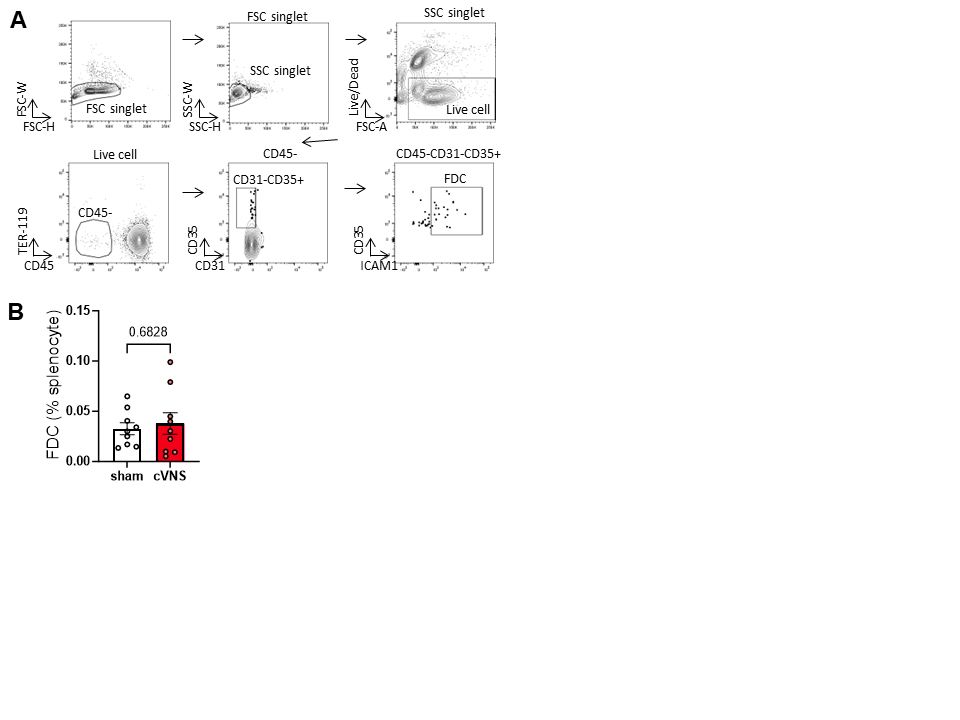

### Figure S3.

**(A)** The gating strategy for FDCs. **(B)** Frequency of FDCs in the spleen of sham and chronic VNS mice. Data are shown as mean +/- SEM with each symbol representing an individual mouse from two independent experiments. Mann-Whitney U test was used to test significance.

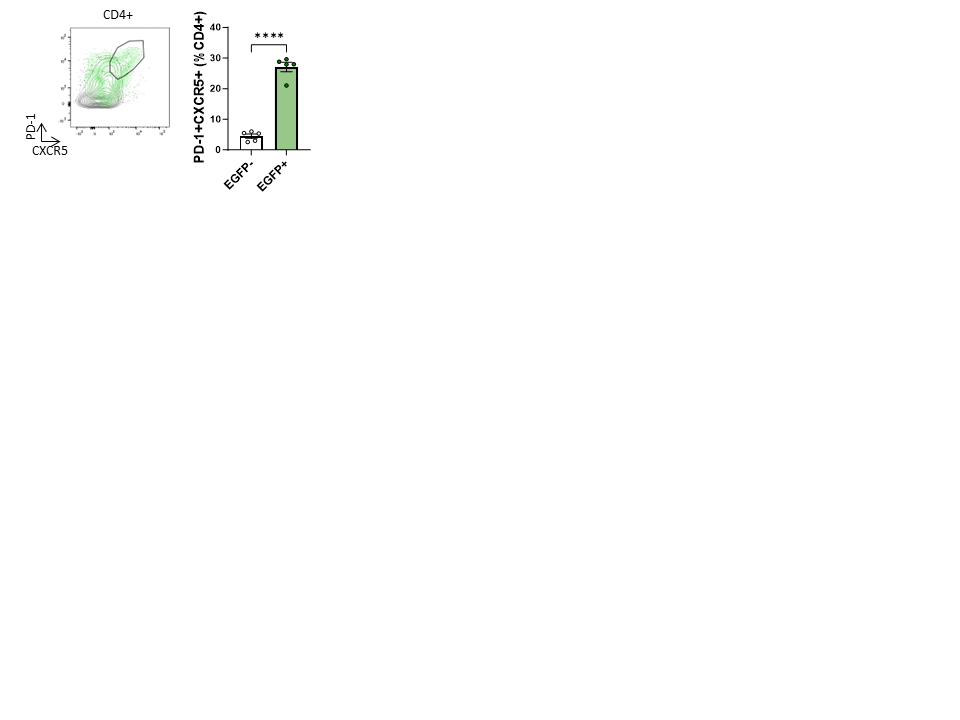

### Figure S4.

Representative plot of PD-1 and CXCR5 expression in eGFP+ (green) and eGFP- (black) CD4+ cells in ChAT^eGFP^ reporter mice. Mice underwent chronic VNS and were analyzed on day 14. Data are shown as mean +/- SEM with each dot representing an individual mouse. Paired t-test was used to test significance.

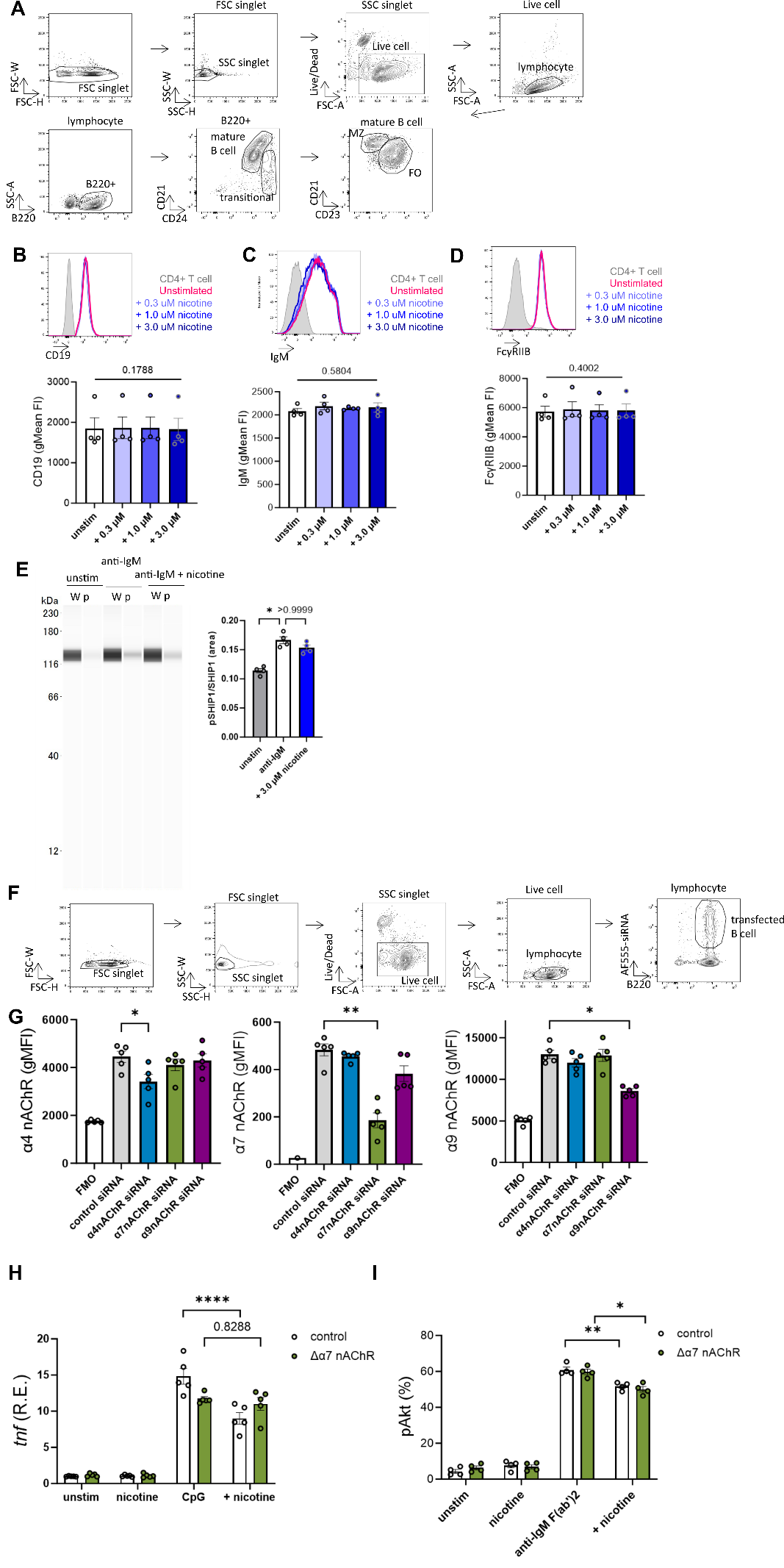

### **Figure S5**.

**(A)** The gating strategy of FO B cells. **(B)** CD19, **(C)** IgM and **(D)** FcγRIIB expression was unaltered by nicotine. FO B cells were incubated with nicotine for 10 min and analyzed by flow cytometry. The expression level on CD4+ T cells is displayed as negative controls. **(E)** SHIP1 phosphorylation unaltered by nicotine. Sorted FO B cells were stimulated with anti-IgM with or without nicotine. Whole SHIP1 (W) and phosphorylated SHIP1 (p) levels were evaluated by western blot. **(F)** The gating strategy of transfected B cells. **(G)** Protein expression evaluation after nAChR knock down. Mean geometric fluorescent intensities were evaluated with flow cytometry. **(H)** Tnf mRNA expression with CpG stimulation in CD19-cre α7nAChR^wt/wt^ (control) and CD19-cre α7nAChR^flox/flox^ (Δα7 nAChR) FO B cells. Sorted FO B cells were stimulated as Fig 5A. **(I)** Akt phosphorylation with anti-IgM stimulation in CD19-cre CHRNA7wt (control) and CD19-cre CHRNA7flox (Δα7 nAChR) FO B cells. Sorted FO B cells were stimulated as Fig 5D. Data are shown as mean +/- SEM. One-way repeated measure ANOVA test with Geisser-Greenhouse correction (B-D), Friedman test with Dunn’s post-hoc comparisons test (E) or paired T test (H, I) was used to test significance.

| **Mouse strains** | | | |
| --- | --- | --- | --- |
| **Abbreviation** | **Strain name** | | **Resource** |
| C57BL/6 | C57BL/6J | | The Jackson Laboratory |
| ChAT^eGFP^ |  | | Dr. Kevin Tracey |
| CD4-cre | B6.Cg-Tg(Cd4-cre)1Cwi/BfluJ | | The Jackson Laboratory |
| ChAT^flox^ | B6;129-Chat^tm1Jrs^/J | | The Jackson Laboratory |
| CD19-cre | B6.129P2(C)-Cd19^tm1(cre)Cgn^/J | | The Jackson Laboratory |
| CHRNA7^flox^ | B6(Cg)-Chrna7^tm1.1Ehs^/YakelJ | | The Jackson Laboratory |
| **Antibodies** | | | |
| **Antigen** | **Fluorochrome** | **Clone/ Catalog No.** | **Supplier** |
| active caspase-3 | AF647 | C92-605 | BD Bioscience |
| B220 | AF488/PerCP/  AF700/BV421 | RA3-6B2 | BioLegend |
| CD3 | APC | 145-2C11 | BD Bioscience |
| CD4 | BV510 | RM4-5 | BioLegend |
| CD19 | unconjugated | EPR23174-145 | Abcam |
| CD21/35 | Biotin | 8C12 | BD Bioscience |
| CD21/35 | PE | 7G6 | BD Bioscience |
| CD23 | PE/Cy7 | B3B4 | Invitrogen |
| CD24 | Pacific Blue | M1/69 | BioLegend |
| CD31 | BV510 | MEC13.3 | BD Bioscience |
| CD45 | PerCP | 30-F11 | BioLegend |
| CD86 | PE/Cy5 | GL-1 | BioLegend |
| CD95 | BV421 | Jo2 | BD Bioscience |
| CXCR4 | BV605 | L276F12 | BioLegend |
| CXCR5 | biotin | 2G8 | BD Bioscience |
| GL7 | PE | GL7 | BioLegend |
| ICAM1 | PE | YN1/1.7.4 | Invitrogen |
| Nicotinic ACh receptor α4 | unconjugated | ab41172 | Abcam |
| Nicotinic ACh receptor α7 | unconjugated  /FITC | ANC-007 | Almone Labs |
| Nicotinic ACh receptor α9 | unconjugated | ab177119 | Abcam |
| NFκB p65 | unconjugated | 8242 | Cell Signaling Technology |
| PD-1 | BV421 | 29F.1A12 | BioLegend |
| Phospho-Akt (pS473) | PE | M89-61 | BD Bioscience |
| Phospho-CD19 (Tyr531) | unconjugated | 3571 | Cell Signaling Technology |
| Phospho-SHIP1 (Tyr1020) | unconjugated | 3941 | Cell Signaling Technology |
| SHIP1 | unconjugated | 2728 | Cell Signaling Technology |
| TER119 | PE eF610 | TER-119 | Invitrogen |
| **Primers** | | | |
| **Gene** | **Assay** | **Assay ID** | **Supplier** |
| Bach2 | Taqman | Mm00464379_m1 | Thermo Fisher Scientific |
| Bcl6 | Taqman | Mm00477633_m1 | Thermo Fisher Scientific |
| Cr2 | Taqman | Mm00801681_m1 | Thermo Fisher Scientific |
| Fcgr2b | Taqman | Mm00438875_m1 | Thermo Fisher Scientific |
| Icam1 | Taqman | Mm00516023_m1 | Thermo Fisher Scientific |
| Ikbkb | Taqman | Mm01222247_m1 | Thermo Fisher Scientific |
| Irf4 | Taqman | Mm00476128_m1 | Thermo Fisher Scientific |
| Pax5 | Taqman | Mm00435501_m1 | Thermo Fisher Scientific |
| Polr2a | Taqman | Mm00839502_m1 | Thermo Fisher Scientific |
| Prdm1 | Taqman | Mm00476128_m1 | Thermo Fisher Scientific |
| Tnf | Taqman | Mm00443258_m1 | Thermo Fisher Scientific |
| Tnfrsf1a | Taqman | Mm00441883_g1 | Thermo Fisher Scientific |
| Vcam1 | Taqman | Mm00516023_m1 | Thermo Fisher Scientific |
| **siRNAs** | | | |
| **Gene** | **Assay ID** | | **Supplier** |
| CHRNA4 | 162556 | | Thermo Fisher Scientific |
| CHRNA7 | 59928 | | Thermo Fisher Scientific |
| CHRNA9 | 502295 | | Thermo Fisher Scientific |

### Table S1. List of mouse strains and reagents used in the study.
